# Genotype Concordance and Polygenic Risk Score Estimation across Consumer Genetic Testing Data

**DOI:** 10.1101/795666

**Authors:** Prag Batra, Kuan-lin Huang

## Abstract

The consumer genomics industry is steadily growing and delivering genetic information to over 10 million individuals. Yet, the implications of using data from different services remain unclear. We investigated the genotyped sites, concordance, and genetic risk estimation using data from three major consumer services. We found low overlap among reported genotyped sites (2.7% across all three). While there was a high concordance between overlapping sites of the two SNP-array services (99.6%), there was a lower concordance between them and a low-pass whole-genome service (73.0%). Finally, we demonstrated that the discrepancy resulted in distinct APOE genotypes and genetic risk scores of Alzheimer’s disease determined using these data. Our results demonstrate genotype results across consumer genomics platforms may lead to different genetic risk estimates, highlighting the need for careful quality control and interpretation.

## INTRODUCTION

The consumer genomics industry is growing rapidly with over 10 million genotyped individuals in 2018.^1^ The services represent the dominant channels to providing genetic data for each individual, yet the implications estimating genetic risk using the resulting data from different companies remain unclear.

## RESULTS

### Different reported genomic sites across three consumer genomics services

We submitted the same saliva sample from an individual at a single time point in each of the three different kits. We then utilized the genotype data provided by each service for further analysis (**Methods**).

We first investigated raw genotype data reported by three major consumer genomics services: 23andMe and AncestryDNA, which used SNP array platforms and a mixed low-pass/imputed whole-genome sequencing platform, Gencove. Low overlap was found among the reported genomic sites (Figure 1A), including 911,444 out of 33,877,548 total sites (2.7%) across all three services and 147,451 out of 1,144,985 sites (12.9%) across 23andMe and AncestryDNA SNP array services. Meanwhile, we also found a substantial number of non-overlapping sites between each of the three services: Gencove, as a low read depth (0.4-1X) and imputed whole-genome sequencing service, offered a high count of 32,732,563 uniquely reported variants while missing 108,221 sites reported by 23andMe and 190,670 sites reported by AncestryDNA. Between 23andMe and AncestryDNA, the non-overlapping sites likely suggest a difference in choice of sites to include on each service’s SNP array. The different reported sites could represent the first source of variability between consumer genomics services that could affect the consistency between downstream analyses.

**Figure 1:**
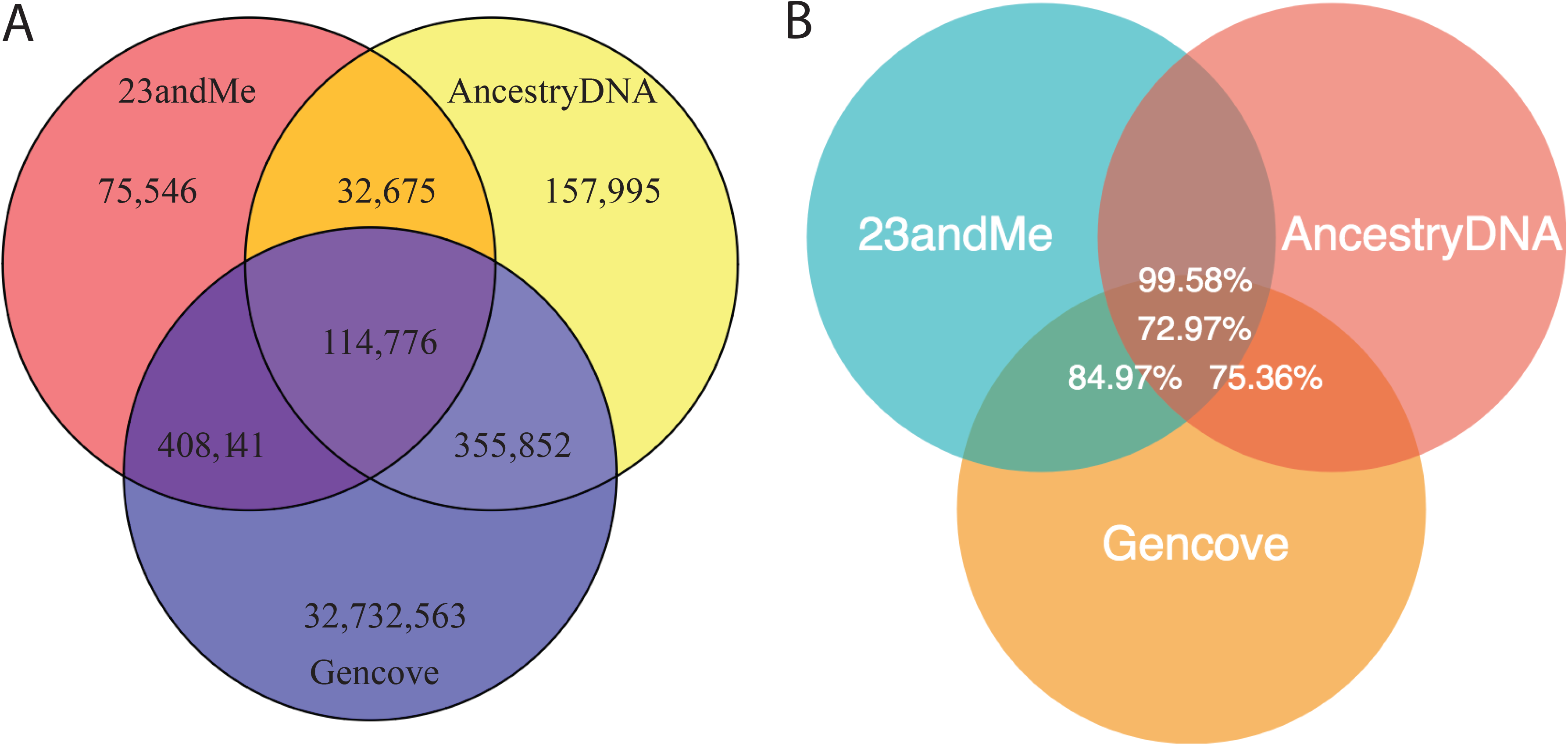
Overlap in site coverage across different consumer genomics services (**A**), and (**B**) the concordance among these overlapping sites.

### Concordance across reported genomic loci

We next investigated whether the genotypes are concordant across sites reported by more than one service. While there was high concordance among overlapping sites between 23andMe and AncestryDNA (99.6%), there was lower concordance between 23andMe and Gencove overlapping sites (85.0%), AncestryDNA and Gencove overlapping sites (75.4%), and sites common to all three services (73.0%, Figure 1B), again suggesting that while Gencove sequencing may cover larger genomic regions over 23andMe and AncestryDNA’s SNP arrays, the low-pass whole genome sequencing could affect the accuracy of the service’s reported variants. The high concordance between 23andMe and AncestryDNA overlapping sites increases confidence in the technical accuracy of each service’s SNP-array technology. Yet, the moderate discordance overall represents a second source of variability that may affect the consistency between services.

### The discrepancy in estimating polygenic risk score (PRS) of Alzheimer’s disease

To evaluate the effect of using data from different consumer genomics companies in assessing disease risk, we applied a polygenic risk score formula to calculate risk to Alzheimer’s disease (AD) (**Methods**). Scoring with the 21 independently-associated SNPs (P < 5E-8) from a recent large-scale AD GWAS of 94,437 individuals,^4^ we found that the genotypes with each of the services amounted to a different score: AncestryDNA = 0.332, 23andMe = 0.525, and Gencove = 0.225. When the missing genotype data of each service was inferred using population minor allele frequency (MAF, **Methods**), as commonly implemented in polygenic risk score algorithms,^3^ the score discrepancy narrowed: AncestryDNA = 0.270, 23andMe = 0.336, and Gencove = 0.225. We noted the limitation that this study’s protocol only collected one individual’s data, thus not allowing for scaling each genetic risk score to a background distribution of all individuals’ data from the same platform.

The effect size from considering each SNP’s genotype showed the sites that contributed to the discordant polygenic risk score estimation (Figure 2). For example, rs429358, one of the SNP specifying APOE genotype, conferred the largest risk effect. However, while 23AndMe and Gencove both identified higher genetic risks given one minor allele of rs429358, AncestryDNA reported two reference alleles without elevated risk. When investigating rs7412, the other allele that together with rs429358 determine the AD APOE ε2/3/4 alleles (**Methods**), AncestryDNA data showed a missing genotype whereas both 23andMe and Gencove showed two reference alleles, confirming the individual as likely APOE ε3/ε4. Another SNP that showed discrepancy was rs6733839, for which 23andMe and AncestryDNA both identified two risk alleles whereas Gencove only identified one. It is also worth noting that the formula for calculating PRS normally utilized population MAF to infer genotypes when there are missing data, and Gencove low-pass WGS provides genotypes at all of the 21 SNPs whereas 23andMe and Ancestry do not.

**Figure 2:**
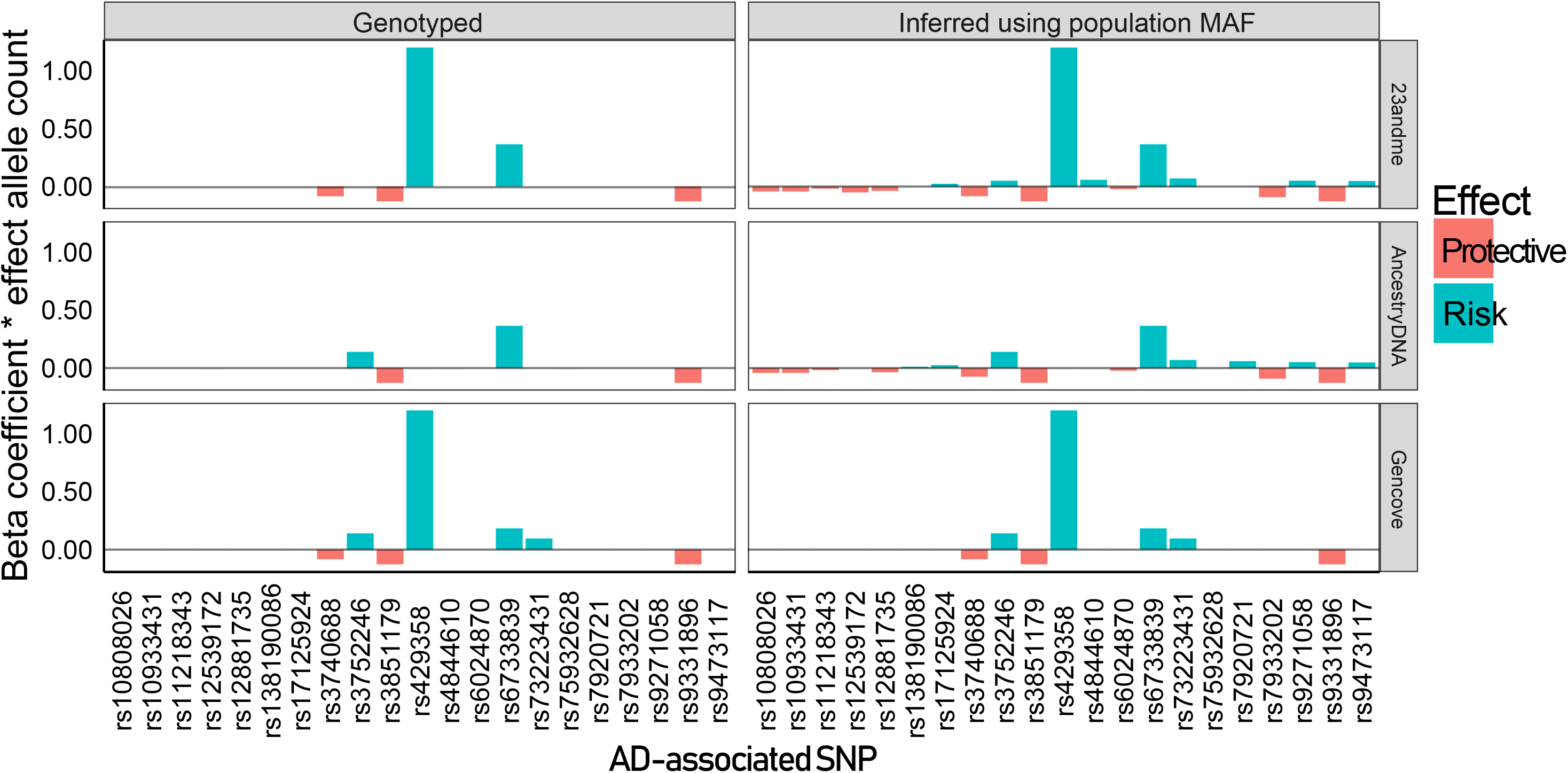
Polygenic risk score (PRS) estimation using 21 independent AD-associated SNPs based on data from 3 different consumer genomic services, including the contribution (beta*effective allele count) of each genotyped SNP (left) or genotyped plus inference using population minor allele frequency (right).

## DISCUSSION

In this study, we have provided a personalized comparison of genotype concordance and genetic risk estimate using consumer genomic data. Our genotype concordance results are consistent with other studies of consumer genetic sequencing services. Our 12.9% site overlap across AncestryDNA and 23andMe is slightly lower than the 16-18% of SNPs reported to be shared across 23andMe, AncestryDNA, and MyHeritage,^7^ as is our 99.6% concordance among overlapping sites between 23andMe and AncestryDNA compared to a 99.6-99.7% concordance reported between Illumina and Affymetrix microarrays and a >99.9% concordance reported between Illumina microarrays.^8^ We note that the genotype concordance with the low-pass NGS results is considerably lower. As sequencing costs continue to drop below the one-thousand-dollar mark,^9^ it is likely that more consumer genetic services will adopt whole-exome or whole-genome sequencing technologies. Given the wide range of different capture kits, sequencing coverages, and variant calling pipelines, considerable variation in DNA-sequencing results has been reported in research and even clinical settings. During the adoption phase of sequencing technology by direct-to-consumer services, this new source of variability from NGS assays needs to be clearly evaluated and communicated.

Our results also demonstrated that analyses using different consumer genomic services may lead to different genetic risk estimation. Given the high general concordance across the genotyping platforms, genetic risk inference based on a single locus is less likely to be biased. However, in this case, we observed a discordant APOE status, the strongest common risk allele contributing to Alzheimer’s disease. In 2017, the FDA approved 23andMe to report APOE status to their customers. In the foreseeable future, one needs to consider the scenario where two services provide conflicting results. Previous studies also reported that differences in the choice of SNPs on microarrays used by different companies can be an important contributor to variability in disease risk^8^ and variability in self-reported phenotypic traits versus genetic testing results.^10^

As direct-to-consumer services begin to infer and report phenotypes, several of which are directly related to consumer health and disease risk,^11^ more sophisticated back-end curation and analysis algorithms, such as polygenic risk scores (PRS), will likely be adopted.^12^ Such reporting introduces additional algorithmic factors as a source of potential variability.

For several complex diseases, recent developments in PRS as determined by multiple disease-associated loci can improve risk stratification in a given population. However, our results demonstrated the potential difficulty to match and compare the genetic risk score provided by different platforms for a single individual. Further, given a polygenic risk score is typically calculated against a “background distribution”, the different customer databases available to each service will also likely contribute to different results. An unbiased standard panel may be designed to benchmark the accuracy of genetic risk estimated by each consumer genomic service. As the use of polygenic risk scores in medicine is being developed,^13^ early discussions for setting up such a standard are required. The algorithms used in each report also deserve clear documentation, transparency, and assessment.

Ancestry reported by each service as of February 25, 2019 was generally concordant regarding major contributors to ancestry, although the services contained important differences regarding minor contributors to ancestry: 23andMe reported 95.1% Chinese; 2.1% Filipino & Austronesian; 0.5% Indonesian, Thai, Khmer & Myanma; AncestryDNA reported 100% China; Gencove reported 91% East Asia and 9% Southeast Asia. This general concordance regarding major contributors to ancestry and variability regarding minor contributors to ancestry is consistent with these other studies showing congruent results for the major contributors to ancestry^7^ despite variability when considering all reported ancestral components.^10^

Several consumer genetics services offer technical white-papers explaining how they infer ancestry using each of their genotype data. For instance, 23andMe uses a three-stage pipeline involving a support vector machine (SVM) classifier, autoregressive pair hidden Markov model, and isotonic regression to compare chromosomal segments to a reference genetic panel of almost 10,000 individuals of homogenous ancestry to identify up to 25 different ancestral components from an individual’s genetic data.^14^ In contrast, while AncestryDNA also compares DNA segments against a reference genetic panel and uses a hidden Markov model, they do not use an SVM classifier or isotonic regression and their panel consists of 16,638 DNA samples across 43 different ethnicities.^15^ While these white-papers represent an attempt to provide transparency, the implications of differences among these algorithms and reference populations may not be fully conveyed or understood by an average consumer, as seen by web blogs containing sporadic consumer confusion over conflicting ancestry results reported by different services. Further, given that the technology and algorithms deployed by each service change over time, the dynamic nature of genetics reports should be accounted for.

Overall, despite the general concordance between these three consumer genetic testing services, the discordant genotype calls and genetic risk score estimates raise the need for enhanced quality evaluation and careful interpretation. Coupled with effective communication of results, DNA data owned by individuals can inform without confusing consumers.

## METHODS

### Comparison of genotypes provided across consumer genomic services

We submitted the same saliva sample from an individual at a single time point in each of the three different kits and then downloaded the genotype and ancestry results reported by each service for further analysis.

To benchmark the technical and reporting implications of multiple genetic testing services, we compared the raw genotype data and ancestry results reported by three major consumer genomics services: 23andMe and AncestryDNA, two of the largest providers using SNP array platforms (Illumina Infinium Global Screening Array^5^ and custom Illumina OmniExpress Plus Genotyping BeadChip,^6^ respectively), and a mixed low-pass (0.4-1X) and imputed (average r^2^ >0.9) whole-genome sequencing platform, Gencove.^2^

As each service provided genotype data in its raw file format, we parsed each file separately to determine common genomic sites and concordance in variant calling at these sites. We retained only variants marked PASS from Gencove’s variant call (VCF) files. Because 23andMe and AncestryDNA reported indels with “I” or “D” and did not provide further descriptions of the exact base changes, we excluded indels from our analysis and just considered single nucleotide polymorphisms (SNPs).

### Determination of APOE genotype

We used two SNPs to determine the APOE genotype in the subject using data from each of the 3 consumer genomics testing kits. The APOE genotype is determined using SNPs rs429358 and rs7412. If both SNPs are T, it is determined as APOE ε2. If they are both C, it is determined as ε4. If rs429358 is a T and rs7412 is a C, it is determined as ε3. If rs429358 is a C and rs7412 is a T, it is determined as ε1.

### Calculation of polygenic risk score (PRS) of Alzheimer’s disease

We used the following standard formula used by algorithms such as PRSice^3^ to calculate PRS, using the combined odds ratio (OR) obtained by Kunkle et al 2019:^4^

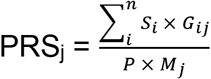

Where PRS is calculated for each consumer genomic service j, the effect size of a disease-associated SNP i is *S*_*i*_, the number of effect alleles is *G*_*ij*_, the ploidy of the sample P is determined as 2 for this individual, and the number of non-missing SNPs among the 21 independently-associated AD SNPs observed for this individual is Mj. We summed up this PRS score using only genotyped SNPs. We also summed up the PRS score by inferring the missing genotypes for SNP i as the population minor allele frequency of the SNP (as reported by Kunkle et al. 2019) multiplied by the ploidy (P = 2) as *G*_*ij*_.

An earlier version of this article was published as a preprint on the bioRxiv server.^16^

## Data Availability

The data that support the findings of this study are available on request from the corresponding author K.H. The data are not publicly available due to the personal genetic data contain information that could compromise research participant privacy/consent.

## Author Contribution

K.H. conceived the study. P.B. and K.H. conducted the analysis. P.B. and K.H. wrote the main manuscript text and P.B. and K.H. prepared the figures and the table.

